# qPCR assays for the detection of two western North American freshwater mussel species, *Anodonta nuttalliana* and *Anodonta oregonensis,* from environmental DNA

**DOI:** 10.1101/372219

**Authors:** Torrey W. Rodgers, Karen E. Mock

## Abstract

We developed species-specific quantitative PCR assays for the detection of two freshwater mussel species native to the western North America, *Anodonta nuttalliana* and *Anodonta oregonensis,* from environmental DNA. These species have experienced dramatic declines over the last century, and are currently threatened in many portions of their range. Improved tools for detecting and monitoring these species are needed. Species-specificity and sensitivity of the assays was empirically tested in the lab, and both assays were also validated with field collected eDNA samples. We found that the assays we designed are species-specific, sensitive, and are effective for detecting *Anodonta nuttalliana* and *Anodonta oregonensis* from environmental DNA samples collected from streams and rivers. These assays will aid in the detection, monitoring, management, and conservation of these threatened species.

## Introduction

The Western United States is home to just five native freshwater mussel species, three of which are from the genus *Anodonta*. Originally, as many as five separate *Anodonta* species were described from the Western US based on morphology, but genetic analyses has shown that only three true genetic lineages exist (Chong et al. 2008). Of these, *Anodonta beringiana* is only known from Alaska and Canada however, the southern extent of its range is uncertain. *Anodonta oregonensis* (synonym *A. Kennerlyi*) ranges from Alaska to Northern California, and *Anodonta nuttalliana* (synonyms *A. californiensis and A. wahlamatensis*) ranges from Washington State to Northern Mexico, and extends east into Idaho, Wyoming, Utah, Nevada and Arizona.

Although not federally listed, native freshwater mussels including *Anodonta* species have experienced dramatic declines in both distribution and abundance in the Western United States due primarily to human impacts. As filter feeders, these species provide important ecosystem services such as improving water clarity, and are a food source for wildlife species. Given ongoing declines, monitoring of native freshwater mussels is clearly important, but traditional sampling requires time-consuming surveys, in some cases necessitating snorkeling or SCUBA diving. Additionally, *Anodonta* species identification requires trained expertise, and in some cases genetic analyses. Thus, improved tools for monitoring these species more efficiently and economically are needed. Environmental DNA (eDNA) is becoming a common tool for aquatic species monitoring, and has been shown to be more sensitive than traditional methods for fish (Wilcox et al. 2016) amphibians (Smart et al. 2015) and reptiles (Hunter et al. 2015). eDNA has also recently been used for detection of freshwater mussels (Stoeckle et al. 2016, Currier et al. 2018, Dysthe et al. 2018). We developed quantitative PCR (qPCR) assays for the detection of two *Anodonta* species, A. *nuttalliana* and *A. oregonensis,* from environmental DNA, and we tested the ability of our assays to detect these species from filtered water samples.

## Methods

### Assay Design

Sequence data for *A. nuttalliana* and *A. oregonensis* spanning the species range from the mitochondrial gene cytochrome oxidase subunit 1 (COI) were compiled from our in-house freshwater mussel sequence database (sequences from 344 samples from 67 populations for *A. nuttalliana* and 95 samples from 23 populations for *A. oregonensis;* supplementary Table S1). Additional sequences from Genbank were also included (27 sequences for *A. nuttalliana* and 11 sequences for *A. oregonensis;* supplementary Table S2). It should be noted that some sequences on Genbank named as *A. oregonensis* are in fact *A. nuttalliana* based on sequence data and clades identified by Chong et al. (2008). For *A. nuttalliana* we included sequences from Washington State (WA; n=25), Oregon (OR; n=112), Idaho (ID; n=12), Utah (UT; n=32), Nevada (NV; n=20), California (CA; n=147), Arizona (AZ; n=9). For *A. oregonensis* we included sequences from samples from OR (n=51), WA (n=31), CA (n=6), British Columbia (n=11) and Alaska (n=3; Figure 1 and Table S1). COI sequence data from non-target, sympatric native species A*. beringiana*, *Gonidea angulata,* and *Margaritifera falcata* were also compiled from our in-house sequence database. COI sequence data from four other potentially sympatric, non-target, non-native taxa: *Corbicula fluminea*, *Dreissena polymorpha, Dreissena bugensis,* and *Lampsilis siliquoidea*, were retrieved from Genbank.

**Figure 1.**
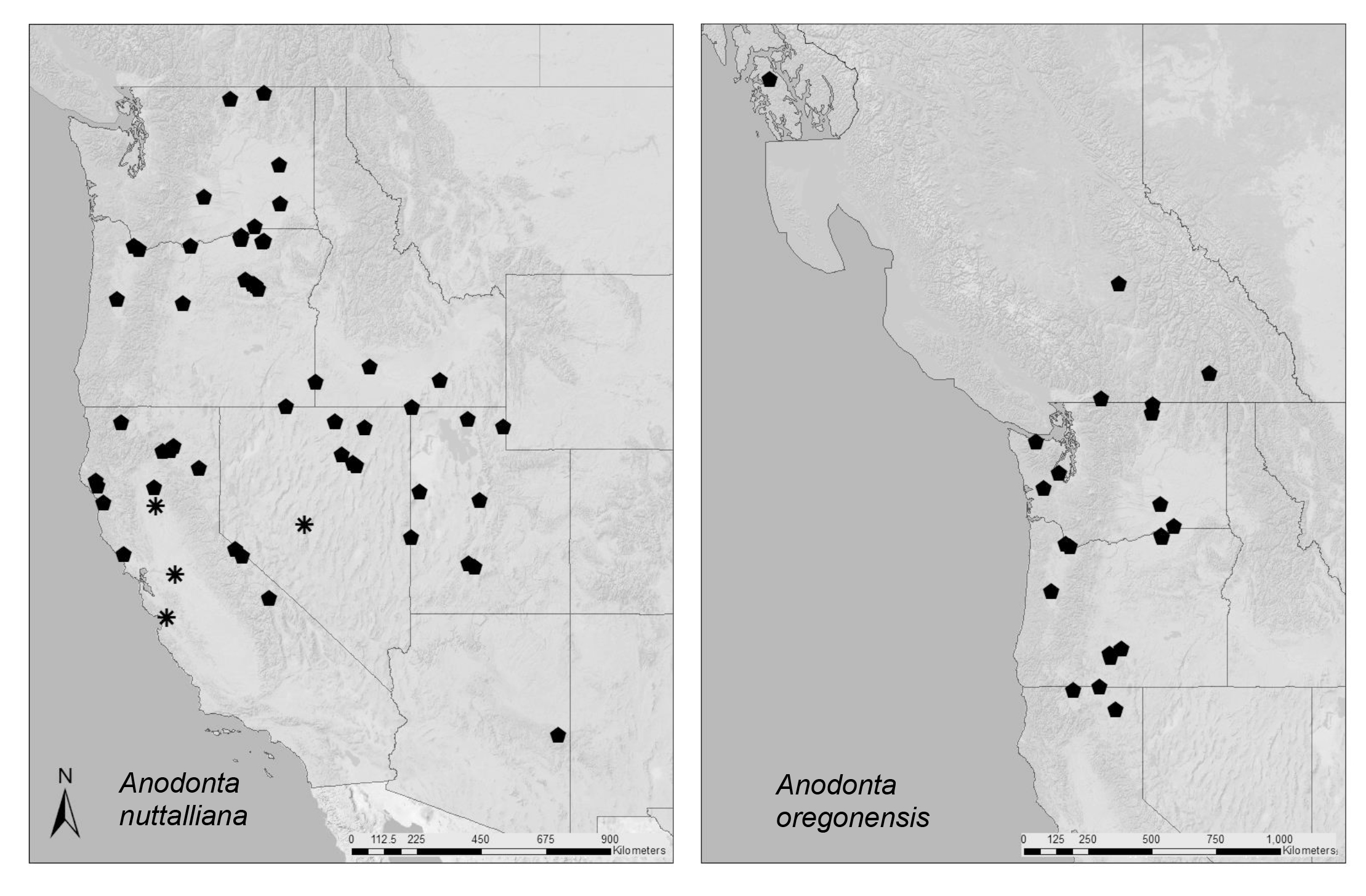
Locations of sample sequences used to design species-specific qPCR assays for the freshwater mussel species *Anodonta nuttalliana* and *Anodonta oregonensis.* Asterisk designates locations were sample sequences possessed a single nucleotide polymorphism with the assay probe AcaCOI2-Pb.

Sequences from all species were aligned using Sequencher software (Gene Codes, Ann Arbor, MI), and we used the online tool DECEPHIR (Wright et al. 2014) to select species specific-primers. We then used ABI primer express software (Applied Biosystems, Foster City, CA) to design a Taqman^®^ Minor Groove Binding qPCR probes and modify primer length to meat melting temperature requirements for qPCR where necessary. For A. nuttalliana we initially selected two candidate primer sets and three Taqman^®^ probes (one probe for the primer set one, and two for primer set two) for testing. For A. oregonensis, we selected one candidate primer set and two Taqman^®^ probes for testing (Table 1).

**Table 1.**
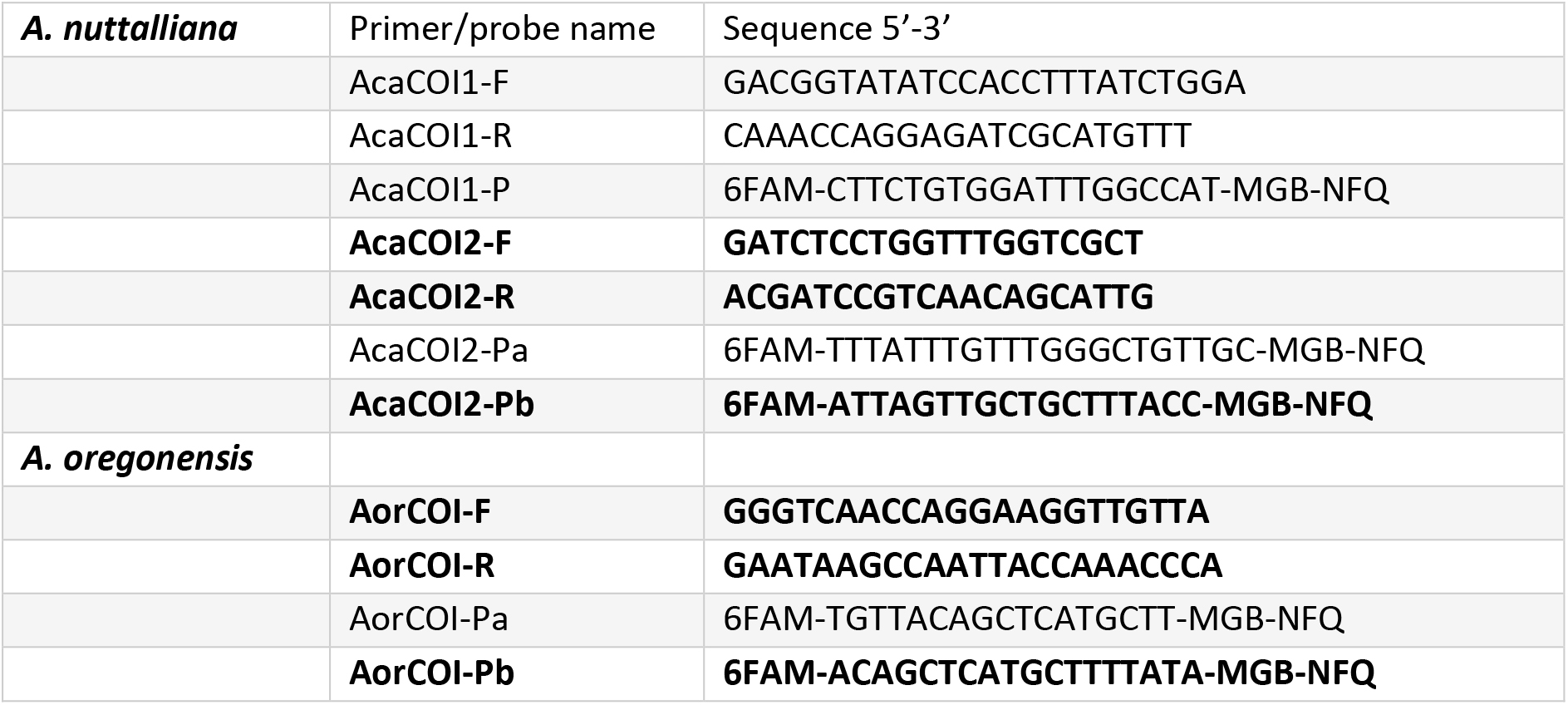
Primer and probe sets developed and tested for detection of *Anodonta nuttalliana* and *Anodonta oregonensis* from environmental DNA. Primers and probes highlighted in bold were chosen as preferable for use in most populations after testing.

### Specificity testing

To test primer specificity in silico, we used NCBI Primer BLAST (Ye et al. 2012) against the NCBI nr database to ensure that no species occurring in the Western United States could potentially amplify and produce false positives.

We initially tested candidate primer sets with SYBR^®^-green qPCR. Test included five target species samples, as well as four samples each of non-target species: *A. nuttalliana/oregonensis*, A*. beringiana*, *G. angulata* and *M. falcata.* qPCR reaction included 7.5 μl SYBR^®^-green power mastermix (Thermo-Fisher) 900 μm of each primer, and 0.1 ng of template DNA in a total reaction volume of 15 μl. Cycling conditions were 95° for 10 minutes followed by 45 cycles of 95° for 15 seconds and 60° for one minute, followed by a melt curve.

We then tested primer sets and probes in Taqman^®^ qPCR. Tests once again included five target species samples and four samples each of non-target species: *A. nuttalliana/oregonensis*, A*. beringiana*, *G. angulata* and *M. falcata.* qPCR reaction included 7.5 μl Taqman^®^ environmental mastermix (Thermo-Fisher) 900 μm of each primer, 250 μm of probe, and 0.1 ng of template DNA in a total reaction volume of 15 μl. Cycling conditions were 95°for 10 minutes followed by 45 cycles of 95° for 15 seconds and 60° for one minute. We ultimately chose one ‘assay’ (a single primer set and probe) for each species based on performance (see methods) for further primer optimization and sensitivity testing. Chosen assays included primer set AnuCOI2 and probe AnuCOI2-Pb for *A. nuttalliana*, and primer set AorCOI and probe AorCOI-Pb for *A. oregonensis* (Table 1).

### Sensitivity testing

To test assay sensitivity, we utilized a MiniGene synthetic plasmid containing the assay sequence (Integrated DNA Technologies, Coralville, IA). The plasmid was suspended in 100 μL of IDTE (10 mM Tris, 0.1 mM EDTA) buffer, linearized by digestion with the enzyme Pvu1, and then purified with a PureLink PCR Micro Kit (Invitrogen, Carlsbad, CA) following standard protocols. The resulting product was quantified on a qubit fluorometer, and quantity was converted to copy number based on molecular weight (Wilcox et al. 2013). For sensitivity testing the product was then diluted to create quantities of 2, 5, 10, 20, 50, and 100 copies/reaction. Each of these quantities was run in six qPCR replicates to determine assay sensitivity.

### Field validation

For *A. nuttalliana* we validated our eDNA assay at 10 total sites in NV (n=6), UT (n=1), OR (n=2), and WA (n=1; Table 2). At all locations live individuals were located either at the time of eDNA sampling (n=8) or within the previous year (n=2). For *A. oregonensis,* our eDNA assay was validated at one site where live individuals had been found in the previous year (Table 2), and also from a tank containing one live individual at the Confederated Tribes of the Umatilla Indian Reservation water lab in Walla Walla, WA. Samples were collected following the protocol outlined in Carim et al. (2016). All samples were run in triplicate qPCR with 4 μl of extraction product, 7.5 μl Taqman^®^ environmental mastermix (Thermo-Fisher) 900 μm of each primer, and 250 μm of probe with cycling conditions described above.

**Table 2.** Sites were eDNA samples were collected to for field validation of qPCR assays for detection of *Anodonta nuttalliana* and *Anodonta oregonensis*.

| Site | State | latitude | longitude | Species detected |
| --- | --- | --- | --- | --- |
| Upper South Fork Humboldt River | NV | 40.63 | -115.74 | *Anodonta nuttalliana* |
| South Fork Humboldt River | NV | 40.69 | -115.83 | *Anodonta nuttalliana* |
| Maggie Creek | NV | 40.89 | -116.18 | *Anodonta nuttalliana* |
| Upper Bruneau River | NV | 41.52 | -115.46 | *Anodonta nuttalliana* |
| Bruneau River | NV | 41.91 | -115.68 | *Anodonta nuttalliana* |
| South Fork Owyhee River | NV | 41.67 | -116.40 | *Anodonta nuttalliana* |
| Salt Creek | UT | 41.68 | 112.23 | *Anodonta nuttalliana* |
| Wildhorse Creek | OR | 45.72 | -118.67 | *Anodonta nuttalliana* |
| Middle Fork John Day | OR | 44.89 | -119.20 | *Anodonta nuttalliana* |
| Walla Walla River | WA | 46.06 | -118.90 | *Anodonta nuttalliana*,<br>*Anodonta oregonensis* |

## Results

### Assay design

For *A. nuttalliana,* we were not able to design a single assay that perfectly matches all haplotypes across the species range due to high levels of haplotype diversity. However, we were able to design assays that perfectly match all sequences in our database from WA, OR, UT, ID, NV and AZ. One individual from one site in NV (Reese River; site ARR from supplementary Table S1) displayed a single SNP (Single Nucleotide Polymorphism) in the probe region of probe AnuCOI2-Pb, however this sequence was a perfect match to probe AnuCOI2-Pa. Thus, use of probe AnuCOI2-Pa may be preferable in this population. A second sample from the Reese River, and all other haplotypes from NV, were a perfect match to both probes. For CA, two samples possessed a single SNP in the reverse primer. A single sample from the San Joaquin River (Site ASV; supplementary Table S1) possessed one SNP in the reverse primer six bp from the 3’ end, but seven other samples collected at the same location were a perfect match to the primer. Likewise, a single sample from the Owens River (Site AOR; supplementary Table S1) also possessed one SNP in the reverse primer 14 bp from the 3’ end, but nine other samples from the same location were a perfect match to the primer. Additionally, all samples from the Sacramento, San Joaquin, and Pajaro rivers in Central California (Sites ASA, ASJ, ASM, ASV, ASQ, and APJ; supplementary table S1) possess a single G-A SNP in the probe region 15 bp from the 3’ end. This SNP appears to be monomorphic in these populations, all 32 samples from these rivers possessed this SNP, so for work in these rivers, a modified probe with a G instead of an A at this position could be used. Due to the polymorphisms in Central California populations described above, if eDNA sampling is conducted in any water bodies outside of those for which sequence data was available for this study (Supplementary Table S1), additional tissue samples from the study location should be collected and sequenced to ensure the optimal eDNA assay is used.

Of the 11 *A. nuttalliana* reference sequences on Genbank from Mexico, five are a perfect match to the assay, whereas the remaining Six possess one SNP 13 bp from the 3’ end of the forward primer, and a second SNP 8 bp from the 3’ end in the reverse primer. Thus if eDNA work is to be conducted in Mexico, a modification of the assay may be warranted (Genbank accession numbers for sequences from Mexico are provided in table S2). For *A. oregonensis*, all haplotypes were a perfect match to the assay.

### Specificity testing

In silico primer specificity checks with NCBI Primer BLAST for *A. nuttalliana* identified just one species with the potential to cross-amplify with our chosen primer set, the congeneric species *Anodonta impura*, which is native to Central Mexico and does not occur in the Western US. For *A. oregonensis* Primer BLAST identified 14 mussel species with the potential to cross-amplify (Supplementary Table S3); however, none occur in the Western United States.

In SYBR^®^-green specificity testing for *A. nuttalliana*, all target samples amplified with a mean Ct of 24.18 (range 22.12-26.33) for primer set AnuCOI1, and mean Ct 23.49 (range 21.52-25.53) for primer set AnuCOI2. All non-target samples amplified at >11 Ct higher than targets for both assays, a range which is suitable for specificity once a probe is added. As primer set AnuCOI2 displayed lower mean Ct values than primer set AnuCOI1, we proceeded with primer set AnuCOI2 for further testing. For *A. oregonensis* all target samples amplified with a mean Ct of 24.98 (range 24.32-26.03). All non-target samples either did not amplify, or amplified at >12 Ct higher than targets, a range which is suitable for specificity once a probe is added. For all primer sets, the melt curve produced a single sharp peak indicating no primer-dimer formation or off target amplification.

In Taqman^®^ qPCR specificity testing for *A. nuttalliana,* all target samples amplified with both probes with a mean Ct of 29.68 (range 28.88-31.24) for probe AnuCOI2-Pa, and mean Ct 28.18 (range 25.50-30.44) for probe AnuCOI2-Pb. No amplification was observed in any non-target samples for either probe. Because probe AnuCOI2-Pb displayed a lower mean Ct value than probe AnuCOI2-Pa, we proceeded with probe AnuCOI2-Pb for sensitivity testing and field validation. For *A. Oregonensis* all target samples amplified with both probes with a mean Ct of 29.31 (range 28.01-31.07) for probe AorCOI-Pa, and mean Ct 29.09 (range 27.62-31.46) for probe AorCOI-Pb. No amplification was observed in any non-target samples for either probe. Because probe AorCOI-Pb displayed a lower mean Ct value than probe AorCOI-Pa, we proceeded with probe AorCOI-Pb for sensitivity testing and field validation.

### Sensitivity testing

For *A. nuttalliana,* all replicates amplified down to five copies per reaction, and 76% (4 of 6) of reactions containing two copies per reaction amplified in qPCR. For *A. oregonensis,* all replicates amplified down to 10 copies per reaction, 50% (3 of 6) of reactions containing five copies per reaction amplified, and 33% (2 of 6) of reactions with two copies per reaction amplified in qPCR.

### Field validation

For both species, eDNA was detected in all field samples collected for that species. For *A. nuttalliana*, All samples collected from NV and UT amplified in all 6 qPCR replicates, except for one low density site (the Upper South Fork of the Humboldt River in NV) in which 2/3 qPCR replicates amplified. In OR, one high density site amplified in all 3/3 qPCR replicates, and at one low density site (middle fork of the John Day) 0/3 replicates amplified in the first round of qPCR, so 6 additional qPCR replicates were run, in which only 1/6 amplified. At the single site from WA, 1/3 qPCR replicates amplified.

For *A. oregonensis*, at the one field site tested, 1/3 qPCR replicates amplified. At this site, no live mussels were encountered by our searches at the time of eDNA sampling, however live mussels had been found the previous year. In the single tank sample tested, 3/3 replicates amplified.

## Discussion

We were able to design and validate sensitive, species-specific qPCR assays for two native freshwater mussel species. These assays performed well for detecting mussels in the field at locations where they occur. Currently, *A. nuttalliana* is listed as vulnerable by the IUCN (Blevins et al. 2016), and some states have recently listed them as species of conservation concern. *A. oregonensis* is currently listed as least concern by the IUCN (Blevins et al. 2016b), however dramatic declines have been observed in some portions of the previous range. Continued monitoring of these species will be essential to document persistence and to locate populations for protection.

Future work should focus on establishing best practices for effectively sampling freshwater mussels with eDNA in the field. For example, research is needed to determine transport distances of eDNA from mussel beds to inform the spatial scale at which samples should be taken from water bodies to ensure populations are not missed by sampling. Research to determine the optimal season to sample for maximum detection probability will also be important. Western freshwater mussels such as *Anodonta* release glochidia into the water column during the spring for dispersal by host fish. Glochida could provide an abundance of eDNA for detection at that time. Alternatively, late summer through winter may be preferable for sampling because high water levels in spring can dilute eDNA, making it harder to detect.

In sum, we believe the eDNA assays described here will serve as a useful tool for future surveys of these species in a more economical and efficient fashion than traditional sampling techniques. This in turn will aid in the detection, monitoring, and conservation of these species across the Western United States.

## Acknowledgements

We would like to thank Krissy Wilson from the Utah Division of Wildlife Resources for seeing the value of using eDNA for western freshwater mussels, and for helping fund this study. We would also like to thank Cynthia Tait, Thomas Franklin, Joseph Dysthe, and Kellie Carim from the United States Forest Service, and Alexa Maine from the Confederated Tribes of the Umatilla Indian Reservation for their help with advice and field work.

